# Differential Contribution of Sensorimotor Cortex and Subthalamic Nucleus to Unimanual and Bimanual Hand Movements

**DOI:** 10.1101/2023.05.23.542003

**Authors:** Christina M. Merrick, Owen N. Doyle, Natali E. Gallegos, Zachary T. Irwin, Joseph W. Olson, Christopher L. Gonzalez, Robert T. Knight, Richard B. Ivry, Harrison C. Walker

**Affiliations:** Department of Psychology, University of California, Berkeley, CA, USA 94720; Department of Bioengineering, University of California, Berkeley, CA, USA 94720; Department of Neurosurgery, University of Alabama at Birmingham, Birmingham, Alabama, USA 35294; Department of Neurology, University of Alabama at Birmingham; Birmingham, Alabama 35294; Helen Wills Neuroscience Institute, University of California, Berkeley, CA, USA 94720; Department of Biomedical Engineering, University of Alabama at Birmingham; Birmingham, Alabama 35294

## Abstract

Why does unilateral subthalamic nucleus deep brain stimulation improve motor function bilaterally? To address this clinical observation, we collected parallel neural recordings from sensorimotor cortex and the subthalamic nucleus during repetitive ipsilateral, contralateral, and bilateral hand movements in patients with Parkinson’s disease undergoing subthalamic nucleus deep brain stimulation. We used a cross-validated electrode-wise encoding model to map EMG data to the neural signals. Electrodes in the subthalamic nucleus encoded movement in a comparable manner for both hands during unimanual and bimanual movements, whereas sensorimotor cortex electrodes displayed a strong contralateral bias. To examine representational overlap in encoding across the two hands, we trained the model with data from one condition (contralateral hand) and used the trained weights to predict neural activity for movements produced with the other hand (ipsilateral hand). Overall, between-hand generalization was poor and this limitation was evident in both SMC and STN. A similar method was used to probe representational overlap across different task contexts (unimanual vs. bimanual). Task context was more important for the STN compared to the SMC indicating that neural activity in the STN showed greater divergence between the unimanual and bimanual conditions. These results indicate that whereas SMC activity is strongly lateralized and relatively context-free, STN integrates contextual information with the ongoing behavior.

**Significance Statement:** Unilateral subthalamic nucleus deep brain stimulation (DBS) improves both contralateral and ipsilateral motor symptoms of Parkinson’s disease. To explore mechanisms for bilateral improvement, parallel neural recordings from the sensorimotor cortex (SMC) and subthalamic nucleus (STN) were recorded in patients with Parkinson’s disease undergoing DBS. Neural activity and muscle activity from the hands were collected while patients performed unimanual and bimanual repetitive hand movements. Activity in SMC primarily encoded contralateral movements and was relatively context-free. In contrast, STN encoded movements in a comparable manner for both hands and was sensitive to the behavioral context.

## Introduction

The basal ganglia and frontal lobe are key nodes in the network supporting voluntary limb movements (DeLong, 1979; Alexander, DeLong & Strick, 1986). Somatomotor regions in the basal ganglia receive inputs from several areas of the cerebral cortex including primary motor cortex (M1), supplementary motor area (SMA), and premotor cortex (PM; Alexander & Crutcher, 1990). Following processing in basal ganglia, movement information returns to these cortical regions via the thalamus (Parent & Hazrati 1995; Middleton & Strick, 2000). In patients with Parkinson’s disease (PD), cell death in the substantia nigra disrupts the cortico-basal ganglia motor loop leading to symptoms such as tremor, rigidity, and bradykinesia (Dauer & Przedborski, 2003). Deep brain stimulation (DBS) of the subthalamic nucleus (STN) or globus pallidus (GP) has not only revolutionized treatment for patients with PD, but has also provided a tool for scientists to further understand PD pathophysiology and basic motor control (Kumar et al., 1998).

Functionally, the basal ganglia have traditionally been implicated in inhibiting or modifying motor plans, processes especially relevant during motor planning and initiation (Mink, 1996). More recent work has highlighted potential contributions to the control of on-going movements (Yttri & Dudman, 2016). In PD patients, STN single unit activity is associated with upper limb movements (Abosch, Hutchison, Saint-Cyr, Dostrovsky & Lozano, 2002; Rodriguez-Oroz et al., 2001) and STN single unit activity and local field potentials (LFPs) can decode grip force (Patil, Carmena, Nicolelis & Turner, 2004; Tan et al., 2016).

Most physiology literature has focused on the contralateral arm relative to the DBS lead. However, various lines of evidence suggest that the STN are engaged during movement with either limb. Bilateral changes in STN oscillatory activity occur during unimanual movements of either hand (Alegre et al., 2005). In addition, phase coherence between the left and right STN increases in the alpha range during unimanual movements, indicating physiological connections between bilateral STN, despite the absence of monosynaptic anatomical connections (Darvas & Hebb, 2014). Perhaps most striking, unilateral implantation of STN DBS improves motor function in both limbs, although the contralateral benefits are larger (Tabbal et al., 2008; Walker, Watts, Guthrie, Wang & Guthrie, 2009).

Here we examine how continuous contralateral and ipsilateral hand movements are encoded in the motor region of the STN and the sensorimotor cortex (SMC). Electrophysiological data were collected during DBS implant surgery from 13 patients, each with a directional DBS lead and an ipsilateral electrocorticography (ECoG) strip over SMC. The participants were asked to produce repetitive voluntary movements with either the ipsilateral or contralateral hand, or with both hands. We employed an electrode-wise encoding model, using the electromyographic (EMG) activity during the task to predict neural activity. We focused on predicting the local motor potential, the time-domain amplitude of the neural time series low-pass filtered at 10 Hz. Prior studies show that this signal correlates well with hand and finger movements (Schalk et al., 2007; Flint, Eric, Jordan, Miller & Slutzky 2013). Our primary goal was to compare hand (contralateral vs ipsilateral) and context (unimanual vs bimanual) encoding in the STN and SMC.

## Method

### Patients

Intracranial recordings were collected from 13 patients (4 women; 59.27 years old). Patients were recruited from University of Alabama at Birmingham (UAB) medical center and represent a subset participants in a randomized, double-blind crossover study of directional versus circular STN DBS for moderately advanced Parkinson’s disease (SUNDIAL trial, clinicaltrial.org: NCT03353688). Inclusion/exclusion criteria for recruitment, screening, enrollment, and DBS surgery were followed strictly. All research procedures were approved by the institutional review boards at UAB, and all patients provided informed consent prior to study participation.

### ECoG strip and DBS lead placement

All surgeries were conducted with the patients awake and “off” dopaminergic medications. Before surgery, pre-op 3T PRISMA brain MR images were co-registered with the intra-op O-arm 2 CT images, and standard frame-based stereotaxy was used to target the STN. A temporary 6 contact Ad-Tech ECoG strip was passed over the “hand knob” of ipsilateral SMC, in the manner pioneered by Starr (De Hemptinne et al., 2013; De Hemptinne et al., 2015; Crowell et al., 2012). We used multi-pass single unit microelectrode recordings, macrostimulation, and intraoperative O-arm 2 CT image to select an appropriate trajectory for the permanent location of the DBS lead. The Boston Scientific Cartesia^TM^ directional lead consists of four rows of electrodes in a 1-3-3-1 configuration, with ring-shaped contacts at the top and bottom rows and two segmented rings with three directional contact segments in each of the middle rows. The leads were positioned at a defined electrophysiological depth, such that the middle directional rows are equidistant from the dorsal STN border based upon the single unit recording profile within that trajectory. Thus, the upper directional row (contacts 5, 6, and 7) is just dorsal to STN in zona incerta / anterior thalamus, and the lower directional row (contacts 2, 3, and 4) is within the dorsolateral sensorimotor STN.

### Behavioral Tasks

A standardized battery of motor behaviors was employed intraoperatively. This included simple, repetitive opening and closing hand movements, variants of Item 3.6 of the Unified Parkinson’s Disease Rating scale (UPDRS). These movements require voluntary flexion-extension at the metacarpophalangeal and proximal interphalangeal joints. There were three movement conditions: contralateral, ipsilateral, and bimanual movements. For the bimanual condition patients were instructed to alternate between the two hands, making an extension/flexion (open/close) movement with one hand and then the other. One participant (P09) was excluded from the bimanual condition because they tended to synchronize the movements of the two hands, making it difficult to differentiate the ipsilateral and contralateral contributions. Each condition was performed for 10 s with the order fixed (contralateral, ipsilateral, bimanual) and a 5-20 s break between blocks. At the start of each block, the verbal instructions “ready, set, go” were provided and the participant was instructed to continue the movement until hearing the word “stop.” An examiner synchronized TTL button presses with each verbal command to mark events in time. All patients completed at least 2 blocks of each condition.

### Data acquisition and preprocessing

Electrophysiological signals were recorded from a BrainVision ActiChamp acquisition system and sampled at 25 kHz without digital filters. We simultaneously recorded LFPs from both the DBS probe and the 6 ECoG contacts over primary sensorimotor cortex. Surface electromyography (EMG) signals were recorded from the hands or arms (bilateral first dorsal interosseous (FDI) muscle, bilateral flexor carpi radialis (FCR) muscle, or both).

### Digital pre-processing

The neural data were low-pass filtered at 500 Hz with a fourth-order Butterworth anti-aliasing filter before down-sampling to 1000 Hz. The data were then high-pass filtered at 0.5 Hz with a third-order Butterworth filter to remove slow drifts in the signal. We re-referenced the signal from each electrode using a common average reference montage within each neural area (i.e., SMC and STN region separately). Electrodes were notch-filtered at 60, 120 and 180 Hz to remove line noise from electronic devices powered by outlets in the operating room. The neural signals were low-pass filtered at 10 Hz to extract local motor potentials (LMPs) which correlate with various aspects of contralateral hand movements (Schalk et al., 2007; Flint, Eric, Jordan, Miller & Slutzky 2013).

Similar to the neural data, EMG signals were low-pass filtered at 500 Hz as an anti-aliasing measure before down-sampling to 1000 Hz. Each EMG channel was z-scored, high-pass filtered at 50 Hz, and full-wave rectified (Flint, Ethier, Oby, Miller & Slutzky, 2012). The EMG data were then low-pass filtered at 10 Hz to render an envelope of movement-related activity. All filters were fourth-order, non-causal Butterworth filters. The EMG data were visually inspected and portions with excessive noise were excluded from further analyses. Blocks that had evidence of movement in the other hand (e.g., contralateral movement during an ipsilateral block) were excluded from the unimanual condition.

### Encoding model

The feature matrix used to predict the neural signal for each electrode consisted of the time-lagged, preprocessed EMG signal from a single hand. Time lags in the matrix extend from 500 ms before movement onset to 500 ms after movement onset 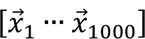. This time range allows compensation for anticipated asynchronies between neural activity and movement, including preparatory neural activity. LMP at each time point, 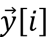, was modeled as a weighted linear combination of the EMG at different time-lags, resulting in a set of regression coefficients, 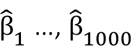,each corresponding to a time lag. To make the regression coefficients scale-free, the EMG features were z-scored before model fitting.

### Model fitting

Regularized (ridge) regression (Hoerl and Kennard, 1970) estimated weights to map EMG signals to the corresponding LMPs for each intracranial electrode.

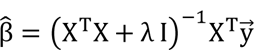

**Equation 1. Ridge regression.** Regression coefficients 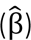 corresponding to each time lag were calculated using the feature matrix of time-lagged EMG data (X) for the LMP signal from each electrode 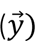. The regularization hyperparameter (λ) was selected based on the inner (validation) test sets. The resulting regression coefficients serve as weights to recombine the time-lagged EMG time series to produce a predicted LMP signal.

For within-arm model fitting, the total dataset consisted of all clean, successful movements performed with either the ipsilateral, contralateral, or bilateral hands (each condition was fit separately). At the outer level, the data were partitioned into five mutually exclusive estimation and test sets. For each test set, the rest of the data acted as the estimation set. For each outer fold, we further partitioned our estimation set into five mutually exclusive inner folds to train the model (80% of estimation set) and predict neural responses across a range of regularization values on the validation set (20% of estimation set). For each inner fold, we tested twenty regularization hyperparameters (λ) evenly spaced on a log scale from 0 to 8, and selected the regularization hyperparameter that provided the best prediction. The averages of the selected regularization parameters and the regression coefficients across the five inner folds were computed to calculate the predicted LMP on the outer test set. This procedure was performed five times at the outer level. Our primary measure is held-out prediction performance (R^2^), quantified as the squared linear correlation between the model prediction and the actual LMP time series, averaged across the five mutually exclusive test sets.

Electrodes were considered predictive (i.e., encoded muscle activity) if they could account for at least 1% of the neural variance (r > .10; R^2^ > .01, see Downey et al., 2020) in the held-out test sets. This criterion was applied separately on predictions derived from contralateral or ipsilateral EMG records. We decided to base our criterion on effect size (R^2^) instead of statistical significance as it is less affected by sample size (which is large when dealing with time series data).

For model fitting across hands, we used the same procedure except that the test set was partitioned from the dataset of the opposite hand. For model fitting across tasks, we used the same procedure with the test set partitioned from the bimanual condition. We partitioned the data in this manner (80% estimation, 20% test) such that fitting procedures for the across hand and across task models were comparable to the within hand models.

## Results

### Predictive electrodes across brain regions

To examine whether muscle activity encoded contralateral and ipsilateral hand movements in individual electrodes over sensorimotor cortex (SMC) and STN, we fit an encoding model to map continuous EMG activity to the LMP signals for 182 electrodes across all patients (n = 13; SMC contacts = 78 and STN contacts = 104). This procedure was performed separately for the two Hand conditions (ipsilateral and contralateral) and two Task conditions (Unimanual and Bimanual) using time-lagged EMG as features in the model. We calculated prediction performance as the square of the linear correlation (R^2^) between the predicted and actual LMP signal, using held-out data from the test set.

Figure 3 summarizes the percentage of predictive electrodes in SMC and STN during either unimanual or bimanual movement. There was a significant difference in the category assignment for the two regions during unimanual movement (χ^2^ = 21.49, *p* < .001). This result is primarily driven by the Contralateral only and Ipsilateral only conditions (Contributions to the χ^2^ value: SMC_contra_ = 6.39, STN_contra_ = 4.80, SMC_ipsi_ = 4.95, STN_ipsi_ = 3.71, SMC_both_ = 0, STN_both_ = 0, SMC_none_ = 0.93, STN_none_ = 0.70; Sharpe (2015). Specifically, electrodes in the SMC were more likely to be predictive of contralateral movement whereas electrodes in the STN were predictive of contralateral and ipsilateral movement to a similar extent.

**Figure 1.**
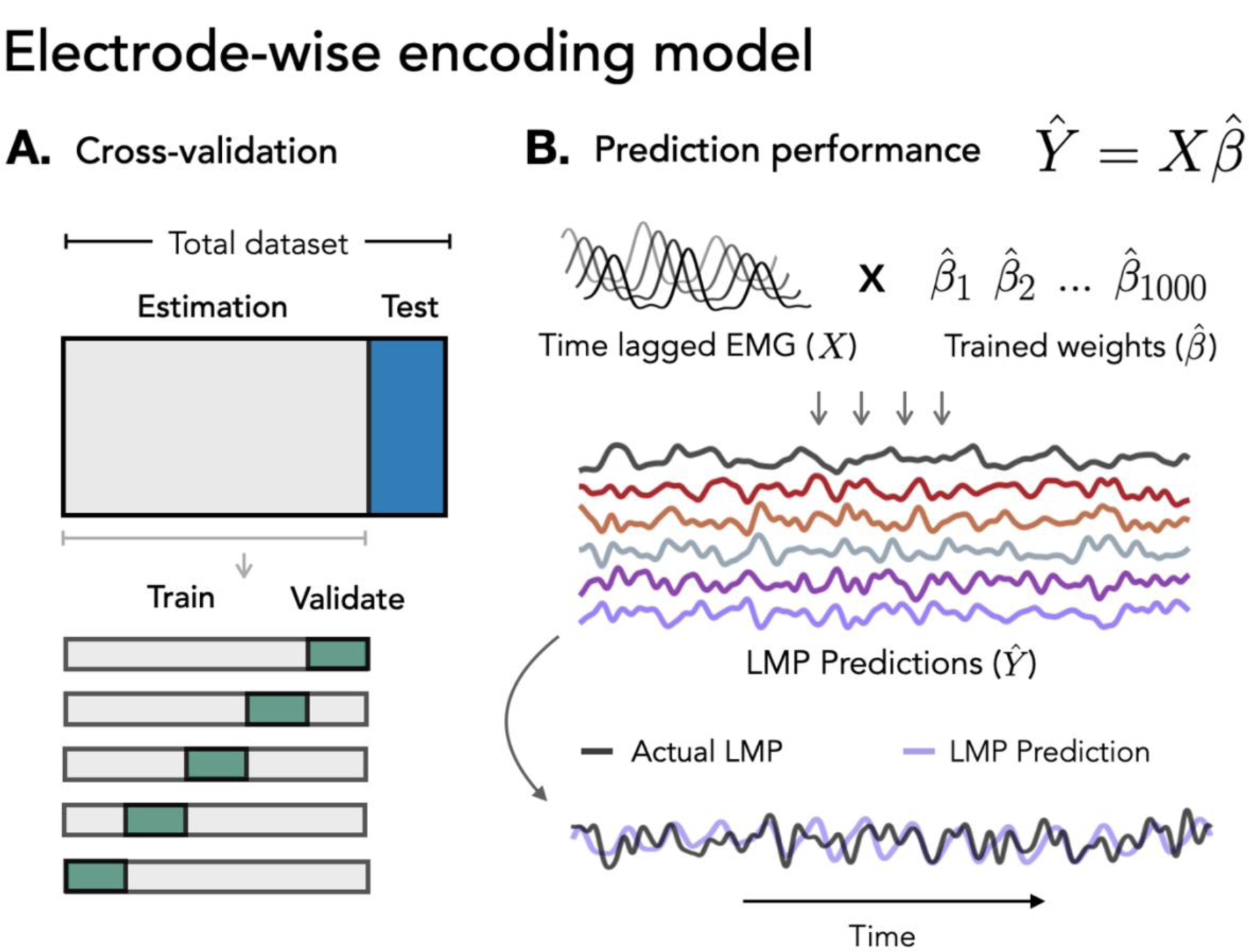
Electrode-wise encoding model. Ridge regression was used to predict neural activity in individual electrodes from continuous EMG activity. **A. Cross-validation.** Nested five-fold cross-validation was used to select the regularization hyperparameter (λ) on inner validation sets. **B. Prediction Performance.** The held-out EMG feature matrix was multiplied with the trained weights to create model predictions.

**Figure 2.**
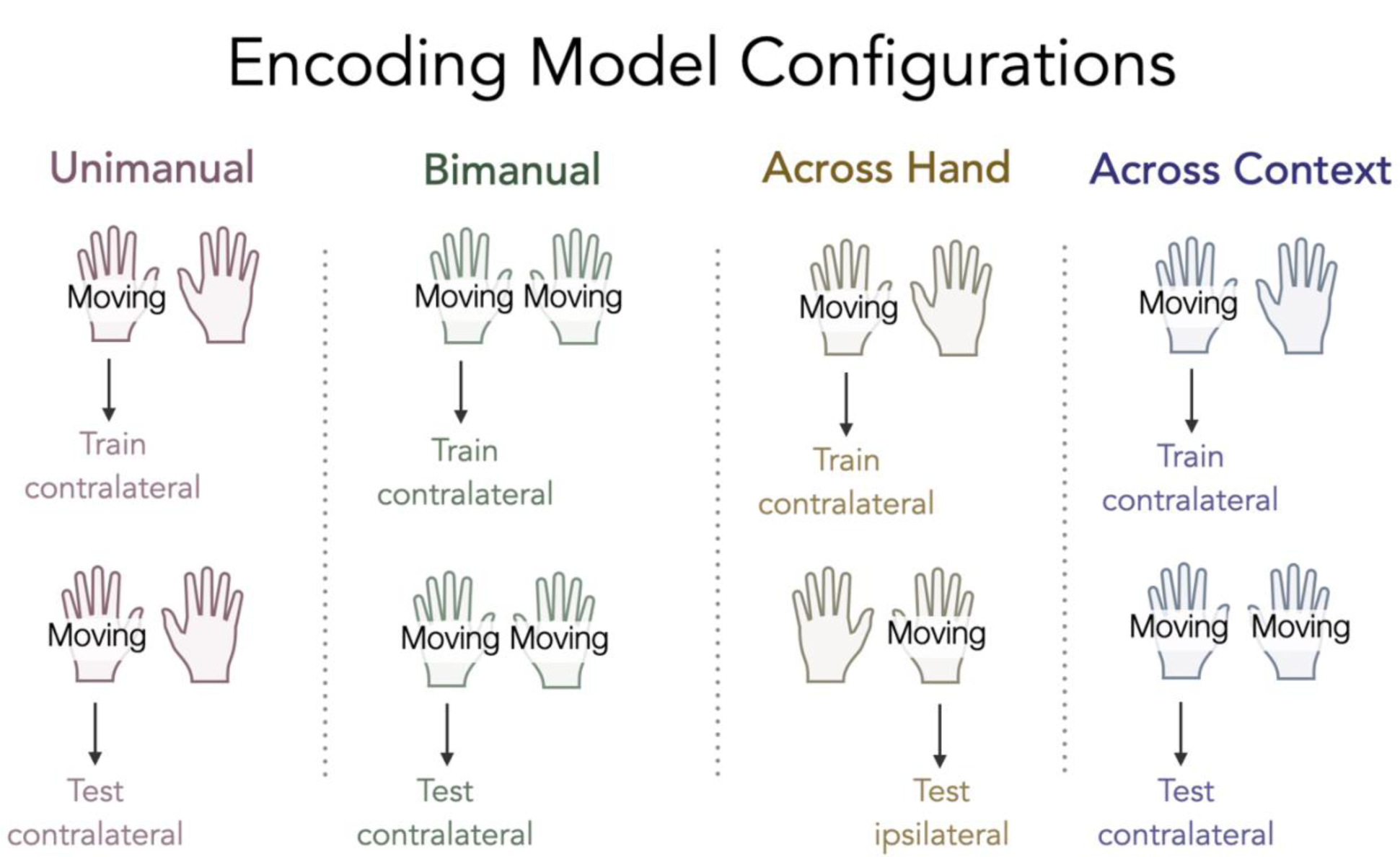
Encoding model configurations. Schematic of the four configurations in our encoding model. For the Unimanual and Bimanual conditions two separate models were created, one examining contralateral movements and one examining ipsilateral movements (ipsilateral not shown for sake of space).

**Figure 3.**
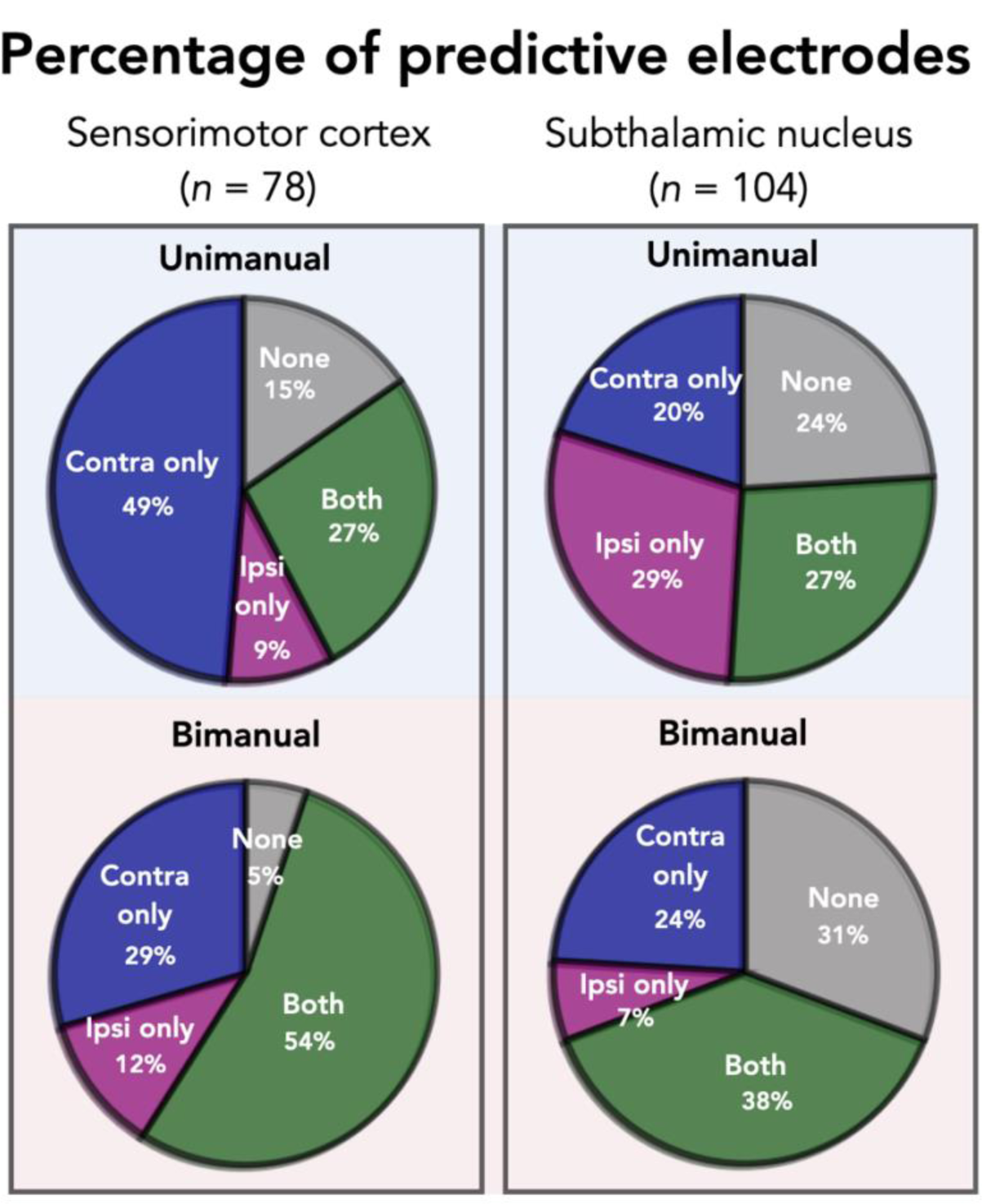
Summary of predictive electrodes across conditions and brain regions. Each electrode was assigned to one of four categories based on predictive performance (R^2^ > .01): Contralateral only; Ipsilateral only; Both; or Neither. During unimanual movement (top row), SMC electrodes are more likely to be predictive of contralateral movement whereas STN electrodes are equally predictive of contralateral and ipsilateral movement. Comparing across brain regions during bimanual movements, significantly more electrodes were classified as None in the STN region compared to SMC.

A similar analysis also revealed a significant difference between STN and SMC in the bimanual condition (χ^2^ = 18.83, *p* < .001). The category with the largest contribution is None, indicating that more electrodes were predictive when both hands are moving in the SMC compared to the STN (Contributions to the χ^2^ value: SMC_contra_ = 0.29, STN_contra_ = 0.22, SMC_ipsi_ = 0.67, STN_ipsi_ = 0.50, SMC_both_ = 1.34, STN_both_ = 1.00, SMC_none_ = 8.47, STN_none_ = 6.35).

### Unimanual prediction performance

Focusing only on the predictive electrodes (excluding electrodes that were categorized as ‘None’ in the prior analysis), we next compared the degree of encoding for contralateral and ipsilateral movement. Figure 4A compares the predictive performance of each electrode during unimanual movement, with the Y-axis plotting contralateral performance and X-axis plotting ipsilateral performance. Values close to the unity line indicate a similar level of prediction performance for each hand; values off the unity line indicate that encoding is stronger for one hand compared to the other. A permutation-based mixed effects model with fixed factors of Hand, Brain Region and a random effect of Patient revealed a significant main effect of Hand (μ_contra_ = .070, μ_ipsi_ = .036, χ^2^ = 16.960, *p* < .001), no effect of Brain Region (μ_SMC_ = .056, μ_STN_ = .050, χ^2^ = .517, *p* > .60), and a significant Hand X Brain Region interaction (χ^2^= 27.221, *p* < .001). The interaction arises from the observation that SMC electrodes more strongly encoded contralateral movement relative to ipsilateral movement (μ_contra_ = .096, μ_ipsi_ = .017, χ^2^ = 37.559, *p* < .001), whereas this bias was not found for the STN electrodes (μ_contra_ = .048, μ_ipsi_ = .052, χ^2^ = 0.139, *p* > .70). Although overall prediction performance did not differ between the two hands for STN electrodes, very few electrodes fell near the unity line, with most electrodes having stronger encoding for either the contralateral or the ipsilateral hand.

**Figure 4.**
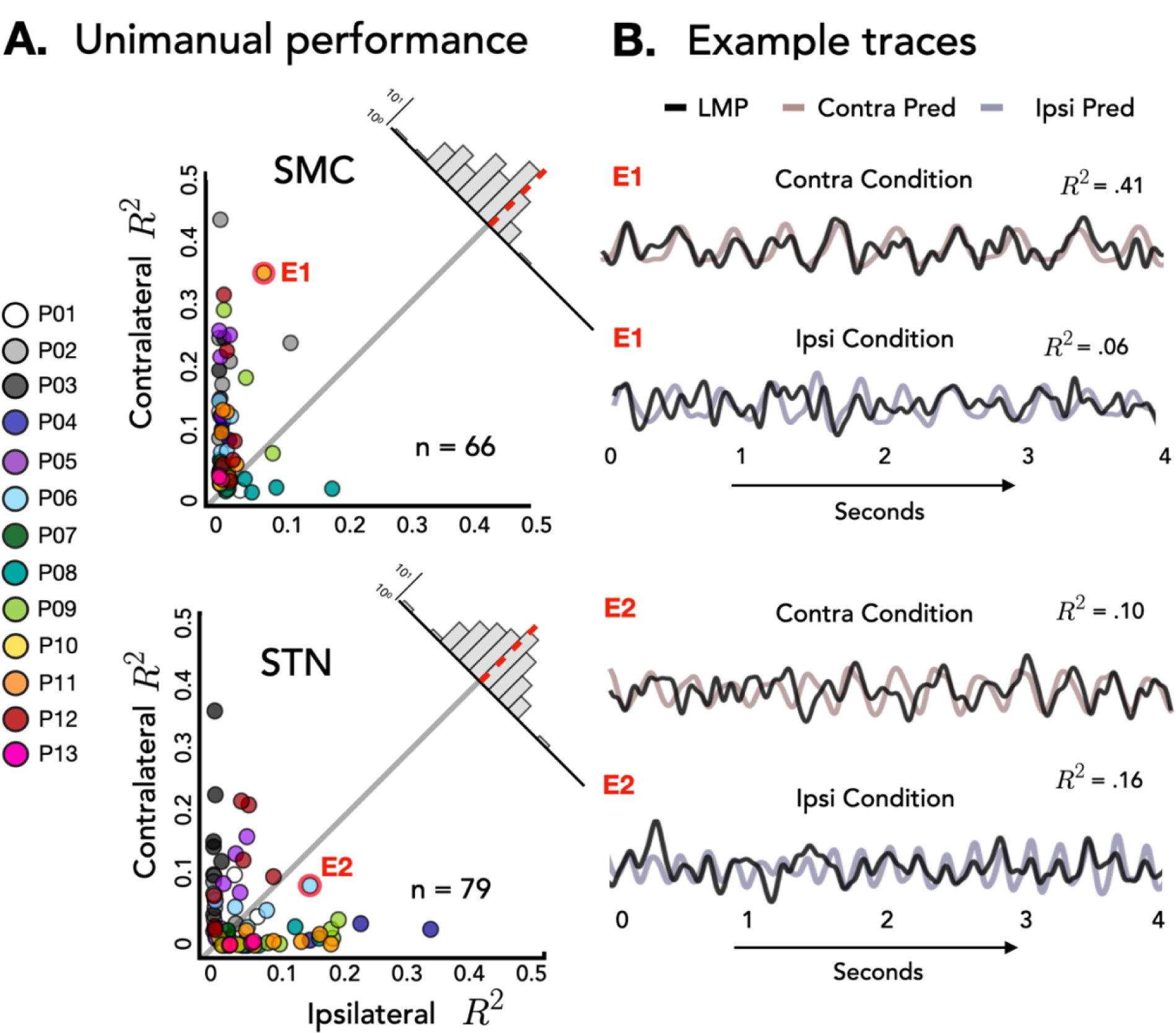
Stronger bilateral encoding in the STN region compared to SMC during unimanual movements. **A. Unimanual prediction performance**. Performance of all predictive electrodes, measured as the square of the Pearson correlation (R^2^), using EMG from either the contralateral (Y-axis) or ipsilateral (X-axis) hand during unimanual movements. Overall performance did not vary by brain regions, but a significant interaction was found with electrodes in the STN region performing equally well across hands whereas the SMC displayed a strong contralateral bias. Data are averaged prediction performance from the five held-out test sets. Upper right corner: Difference distribution for each brain region. **B. Example traces.** Held-out predictions of the LMP time series for one test set of E1 and E2 during unimanual movements.

We next compared prediction performance across the two brain regions for each hand given that we had a significant Hand X Brain Region interaction. Although we did not find a main effect of Brain Region, the simple effects showed that SMC electrodes performed better than STN electrodes during contralateral movement (μ_SMC_ = .096, μ_STN_ = .048, χ^2^= 15.133 *p* < .001), whereas STN electrodes performed better than SMC electrodes during ipsilateral movement (μ_SMC_ = .017, μ_STN_ = .052, χ^2^ = 20.751 *p* < .001).

### Bimanual prediction performance

We next examined if differential contralateral and ipsilateral encoding in SMC and STN is also observed during bimanual movements. Figure 5A compares the predictive performance of each electrode during bimanual movement when the prediction was based on the EMG features from either the contralateral (Y-axis) or ipsilateral (X-axis) hand. There was a significant main effect of Hand (μ_contra_ = .077, μ_ipsi_ = .041, χ^2^ = 13.356, *p* < .001), Brain Region (μ_SMC_ = .075, μ_STN_ = .046, χ^2^ = 7.781, *p* < .05), and interaction of these factors (χ^2^ = 8.899, *p* < .01). The interaction was similar to that observed in the unimanual analysis: Electrodes over SMC encoded contralateral movement better than ipsilateral movement (μ_contra_ = .108, μ_ipsi_ = .042, χ^2^= 21.359, *p* < .001) whereas electrodes in the STN showed similar encoding for contralateral and ipsilateral movement (μ_contra_ = .051, μ_ipsi_ = .040, χ^2^= 1.103 p > .30). In comparison to the unimanual condition (Fig 4A, STN region), in the bimanual condition (Fig 5A, STN region) more electrodes fell near the unity line suggesting these electrodes encode both hands to a similar degree. The main effect of Brain Region in the bimanual condition arose because predictive performance was better overall in SMC than STN in the bimanual condition, a result that we did not observe in the unimanual conditions.

**Figure 5.**
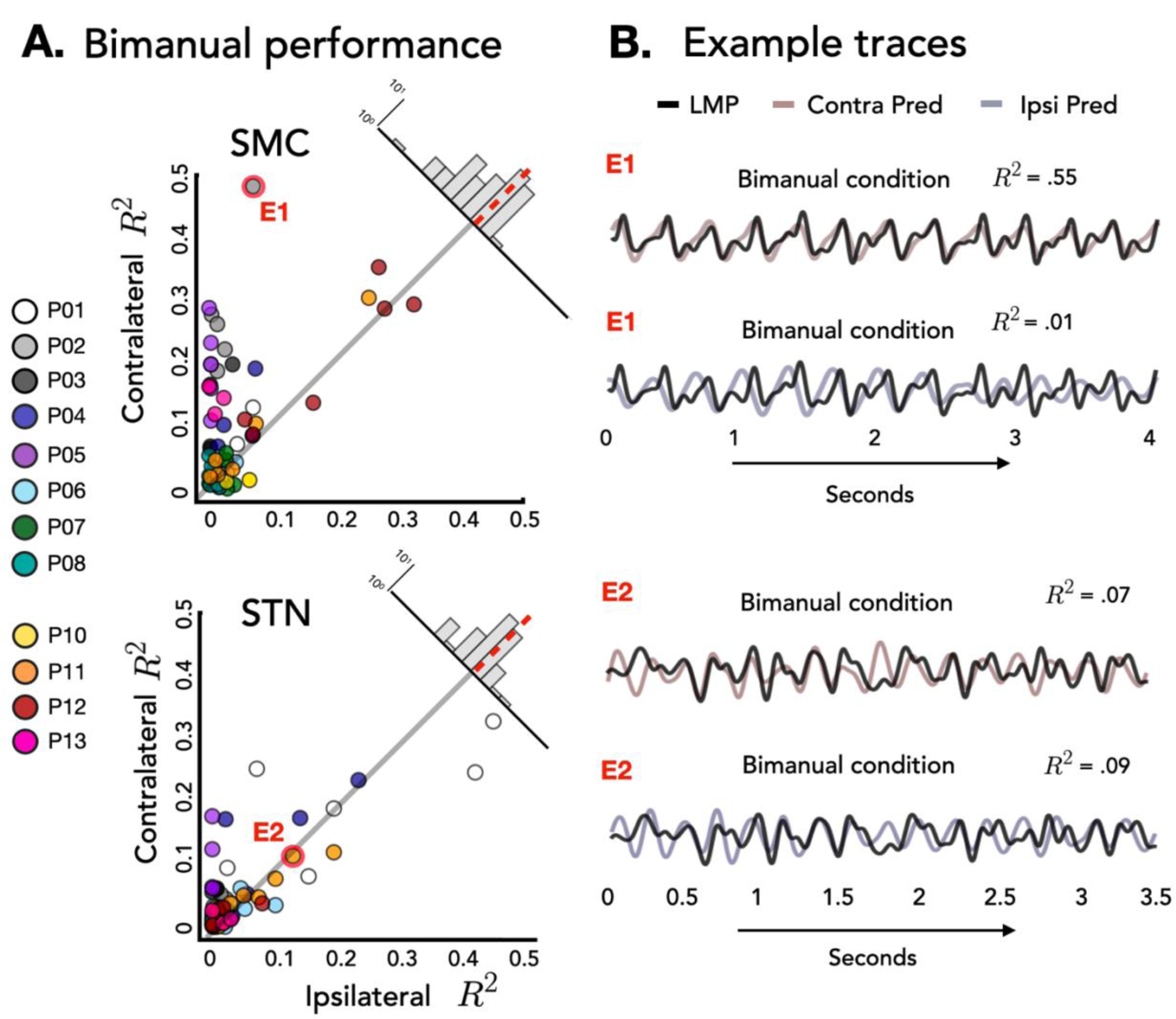
Stronger bilateral encoding in the STN region compared to SMC during bimanual movements. **A. Bimanual prediction performance.** Performance of all predictive electrodes, measured as the square of the Pearson correlation (R^2^), using EMG from either the contralateral (Y-axis) or ipsilateral (X-axis) hand when both hands were moving. A significant interaction was found between hand and brain region with electrodes in the STN region performing equally well across hands whereas a strong contralateral bias was found in SMC. Data are averaged prediction performance from the five held-out test sets. Upper right corner: Difference distribution for each brain region. Insets: Time-locked average of LMP for contralateral and ipsilateral hand opening for E1 and E2. **B. Example traces.** Held-out predictions of the LMP time series for one test set of E1 and E2 when both hands were moving.

To statistically compare predictive performance between the unimanual and bimanual conditions, we fit a permutation-based mixed effects model with three factors, Hand, Brain Region, and Task, along with the random effect of Patient. The two-way interactions involving the new factor Task were not significant (Task X Hand: χ^2^ = 0.012 *p* > .90; Task X Brain Region: χ^2^ = 3.162 *p* > .10), but there was a significant three-way interaction (χ^2^ = 36.470 *p* < .001). Analyzing simple effects, ipsilateral encoding increased during bimanual movement in SMC (μ_uni_ = .017, μ_both_ = .042, χ^2^ = 8.526 *p* < .001; rightward shift of SMC data points in Fig 5A relative to 4A), but remained approximately the same in the STN (μ_uni_ = .051, μ_both_ = .041, χ^2^ = 0.518 *p* > .40). Contralateral encoding did not change across task condition in either region (all χ^2^’s < 0.751 *p*’s > .40). Thus, ipsilateral encoding increased in SMC but not STN when both hands were moving, although it remained much weaker overall in SMC.

### Across hand generalization

The preceding analyses focused on encoding within-hand predictions, with the predictions being generated using held-out data from the same conditions that has been used to derive the model. To examine the degree of overlap in neural representations for contralateral and ipsilateral movement, we examined across hand prediction performance. To this end, we trained the encoding model with the EMG data from one hand and used the trained weights to create predictions for the EMG data from the other hand. Figure 6A shows the percent change from within hand predictions to across hand predictions for all electrodes. Values close to zero indicate good across hand generalization; large negative values indicate poor across hand generalization. We found poor across hand generalization for both brain regions shown by the large decrement in prediction performance (μ_SMC_ = −91%, μ_STN_ = - 86%). Using a permutation-based mixed effects model with a fixed factor of Brain Region and a random effect of Patient, we found no difference in hand generalization (quantified as percent change) between the SMC and the STN (χ^2^ = 1.54 *p* > .20).

**Figure 6.**
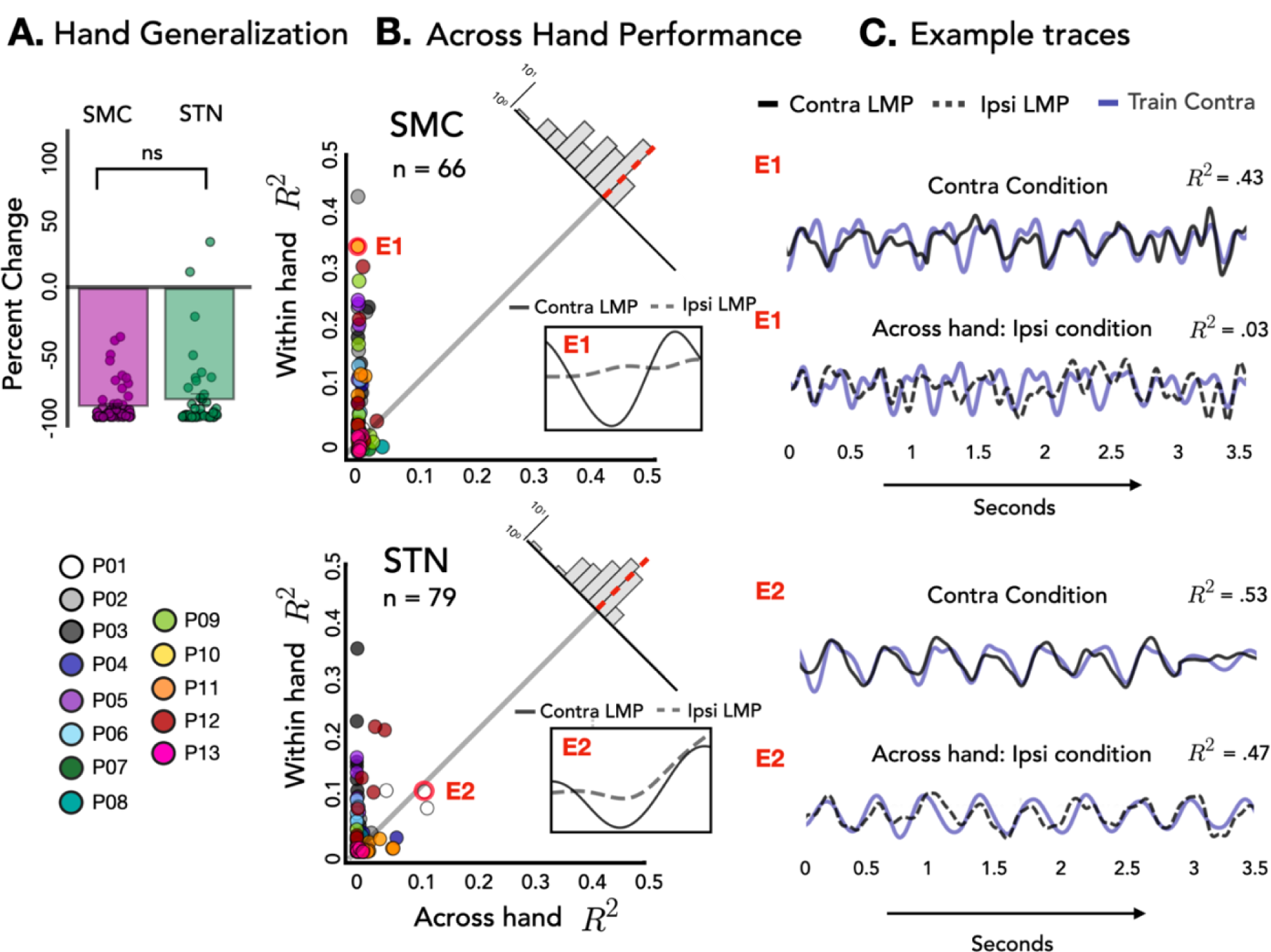
Poor across hand generalization in SMC and STN. A. Hand generalization. Percent change from within hand R^2^ to across hand R^2^ was calculated for each electrode for SMC (magenta) and STN (green). No difference in hand generalization was found between the two brain regions. **B. Across hand performance.** Performance of all predictive electrodes, measured as the square of the Pearson correlation (R^2^), using EMG from either the same hand (Y-axis) or training and testing across hands (X-axis) during unimanual movements. Data are averaged prediction performance from the five held-out test sets. Upper right corner: Difference distribution for each brain region. Insets: Time-locked average of LMP for contralateral and ipsilateral hand opening for two electrodes (E1 and E2). **C. Example traces.** Held-out predictions of the LMP time series for one test set of E1 and E2, the first being within hand and the second being across hand predictions.

Figure 6B shows the predictive performance of each electrode when the model was trained and tested on the same hand (train contralateral, test contralateral; Y-axis) versus across hands (train contralateral, test ipsilateral; X-axis). Values close to the unity line have more overlapping neural representations during contra- and ipsilateral movement, whereas electrodes off the unity line encode the two hands differentially. Figure 6C shows within and across hand predictions for one electrode in SMC and one electrode in STN. We note that although there was not a significant difference in hand generalization across the two areas, a small subset of electrodes generalized well across hands in the STN region (i.e., E2 in Fig 5).

### Context generalization

We next examined context generalizability, asking if encoding changes between the unimanual and bimanual conditions. To this end we trained the encoding model with the EMG data from the contralateral hand in the unimanual condition and used those weights to predict neural activity during movements in the bimanual condition (Figure 7). Figure 7A shows the percent change for each electrode from within context predictions to across context predictions. Values close to zero indicate good across context generalization; large negative values indicate poor across context generalization.

**Figure 7.**
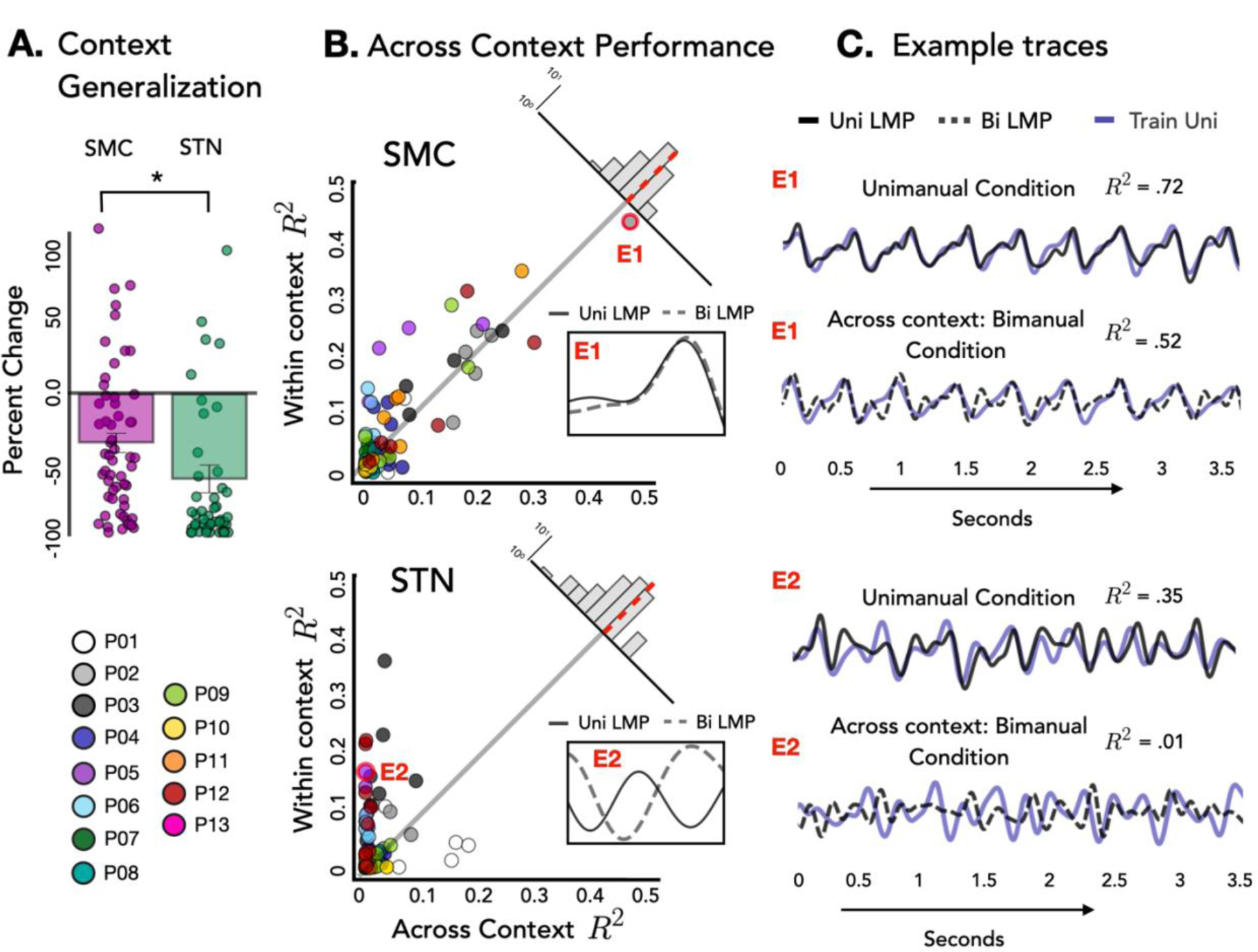
Across context prediction performance. **A. Context generalization.** Percent change from within task R^2^ to across task R^2^ was calculated for each electrode for SMC (magenta) and STN (green). A significant difference was found in context generalization between the two brain regions, with electrodes within the SMC showing stronger context generalization. **B. Across Context Performance.** Performance of all predictive electrodes, measured as the square of the Pearson correlation (R^2^), using EMG from either the same task (unimanual; Y-axis) or training and testing across tasks (unimanual to bimanual; X-axis). Data are averaged prediction performance from the five held-out test sets. Upper right corner: Difference distribution for each brain region. Insets: Time-locked average of LMP for contralateral and ipsilateral hand opening for E1 and E2. **C. Held-out predictions.** Held-out predictions of the LMP time series for one test set of E1 and E2 when both hands were moving.

To assess context generalization, we fit a permutation based mixed effects model with a fixed factor of Brain Region and a random effect of Patient. Across task generalization was greater in the SMC compared to the STN (μ_SMC_ = −35%, μ_STN_ = −68%, χ^2^ = 10.10 *p* < .005), indicating that the STN is more sensitive to context. Specifically, in the STN we saw a larger decrement in across task generalization; thus, the profile of contralateral neural activity differs for contralateral movement produced alone (unilateral) or concurrently with ipsilateral movement (bimanual). In contrast, the profile of contralateral neural activity in SMC remained more similar across unimanual and bimanual conditions. Figure 7B shows the predictive performance of each electrode when the model was trained and tested within the same condition (train unimanual contra, test unimanual contra; Y-axis) or across conditions (train unimanual contra, test both hands contra; X-axis).

## Discussion

Most DBS studies examine physiological and outcome measures focused on the contralateral arm relative to the implanted lead. Yet bilateral changes in STN oscillatory activity are observed during unimanual movement and unilateral implantation of STN DBS improves motor function in both limbs (Alegre et al., 2005; Tabbbal et al., 2008; Walker, Watts, Guthrie, Wang & Guthrie, 2009). Motivated by these observations, we collected electrophysiological data in the STN and the SMC during contralateral and ipsilateral movement. We fit an encoding model to map continuous EMG activity to neural activity and examined the extent of movement encoding in STN and SMC. In addition to the standard within-condition analyses, we also asked how well the encoding model could generalize across hands and across context. By using identical features (the EMG signals) to simultaneously fit SMC and STN field potentials, we avoid some potential confounds (e.g., motor noise) that may limit comparisons across the two brain regions.

We observed differential encoding patterns in STN and SMC during unimanual and bimanual movements. There was a strong contralateral bias in the SMC, whereas in the STN both hands were encoded equally well. However, there was little representational overlap across hands and the degree of generalization did not differ for the two brain regions. This result suggests that although the STN is encoding both hands to a similar extent, the temporal pattern of activation differs for ipsilateral and contralateral movements. We also found that STN had weaker across context generalization compared to SMC. Taken together, these data suggest that activity in the SMC is strongly associated with contralateral movement independent of context whereas STN activity is bilateral and more context dependent.

### Movement encoding in the SMC and STN

The current results provide further evidence that STN activity is associated with movement kinematics (Yttri & Dudman, 2016), and overall, the level of encoding in STN was comparable to that observed in SMC during unimanual movement.

Interestingly, encoding in SMC and STN diverged depending on the hand used for encoding: For contralateral movement, SMC outperformed STN in predicting neural activity whereas for ipsilateral movement, STN outperformed SMC.

Examining motor encoding during bimanual movement, we found that SMC had higher levels of encoding compared to STN, a difference that was not found during unimanual movements. In terms of the categorical analysis, the number of electrodes that encoded movement in the SMC increased from unimanual to bimanual movement. This trend was not found in the STN, and descriptively the number of electrodes encoding movement decreased from unimanual to bimanual movements. Despite this increase in the SMC, the motor encoding remained strongly contralateral whereas in the STN, more electrodes showed bilateral encoding. The SMC pattern may arise due to the increase in activity in the opposite hemisphere during bimanual movement, signals that would be communicated across the corpus callosum via direct connections between homologous regions or indirectly via communication between secondary motor and association cortices (Kazennikov, Hyland, Corboz, Babalian, Rouiller, & Wiesendanger, 1999). This may require a concomitant increase in contralateral encoding to ensure the coordinated movement of that hand.

This speculation points to one limitation of the current study: Our bimanual task required little coordination between the two hands. Future research could use encoding models to track neural activity during a behavioral task that requires coordination of both hands towards a common goal (e.g., opening a jar). Such tasks would also prove beneficial in that the movements of the two hands would be quite distinct, making it easier to separate the individual contributions of the ipsilateral and contralateral hands during bimanual movements.

### Task Generalization

The STN has connections with several cortical (PFC, PMC, SMA & M1) and non-cortical regions (Thalamus, Globus Pallidus, Cerebellum; Benarroch 2008), making it an ideal hub to integrate motor and non-motor information. The STN is hypothesized to integrate environmental cues with ongoing behaviors (Sauleau et al., 2009; Péron, Frühholz, Vérin & Grandjean 2013). Consistent with this idea, we found that context (i.e., task condition) was more important in the STN compared to the SMC. Specifically, the temporal pattern of neural activity tracking contralateral movement in the STN changed when the ipsilateral hand was also engaged in the task. In contrast, neural activity tracking contralateral movements in the SMC was similar across the unimanual and bimanual conditions. This result is in line with the idea that neural activity in the SMC is strongly associated with movement of the contralateral limb, with effector independent activity becoming more evident in premotor and posterior parietal regions (Merrick et al., 2022).

### Implications for adaptive DBS

While DBS has been used as a therapeutic device to treat Parkinson’s disease for 30 years, the majority of DBS applications involve constant-amplitude neurostimulation (Lozano et al., 2019). Adaptive DBS (aDBS) has the potential to adjust stimulation parameters in response to electrophysiological biomarkers, with tremendous potential for improving patient outcomes (Gilron et al., 2021; Starr 2018). Understanding the functional role of the STN is likely to be important for developing optimal aDBS algorithms. Our results show that the STN has similar encoding capability for contralateral and ipsilateral movement and is context sensitive. This suggests that optimizing an aDBS device based on symptoms on both sides of the body could outperform algorithms that focus solely on the contralateral side. This perspective is further supported by the observation that unilateral DBS improves symptoms on the ipsilateral side of the body, although to a lesser extent than the contralateral side (Walker, Watts, Guthrie, Wang & Guthrie, 2009). Further studies should examine the degree of bilateral encoding in the STN in other task domains such as upper arm movements or foot movements.

## Acknowledgments

This research was supported by the National Institutes of Health: R01-NS097480, R35-NS116883, UH3-NS100553 and R01-NS119520.

